# ORN: Extracting Latent Pathway Activities in Cancer with OR-gate Network

**DOI:** 10.1101/2020.06.07.137992

**Authors:** Lifan Liang, Kunju Zhu, Songjian Lu

**Author notes:** Songjian Lu, Department of Biomedical Informatics, University of Pittsburgh, Pittsburgh, PA, 15206, United States.

## Abstract

Pathway level understanding of cancer plays a key role in precision oncology. In this study, we developed a novel data-driven model, called the OR-gate Network (ORN), to simultaneously infer functional relationships among mutations, patient-specific pathway activities, and gene co-expression. In principle, logical OR gates agree with mutual exclusivity patterns in somatic mutations and bicluster patterns in transcriptomic profiles. In a trained ORN, the differential expression profiles of tumours can be explained by somatic mutations perturbing signalling pathways. We applied ORN to lower grade glioma (LLG) samples in TCGA and breast cancer samples from METABRIC. Both datasets have shown pathway patterns related to immune response and cell cycles. In LLG samples, ORN identified multiple metabolic pathways closely related to glioma development and revealed two pathways closely related to patient survival. Additional results from the METABRIC datasets showed that ORN could characterize key mechanisms of cancer and connect them to less studied somatic mutations (e.g., BAP1, MIR604, MICAL3, and telomere activities), which may generate novel hypothesis for targeted therapy.

## INTRODUCTION

In the past decades, the emergence of high-throughput technology has greatly facilitated cancer research. One major insight from these efforts is that although somatic genomic alterations (SGA) vary from patient to patient, they exhibit similar patterns on pathway level (1). That is, different somatic mutations in different samples perturb the same signalling pathway and drive similar tumour phenotype. Based on this insight, many anti-cancer therapies targeting certain pathways have been successfully deployed (2). Unfortunately, most of these therapies are only effective for specific subpopulations. And patient responsiveness is often difficult to predict. For example, despite the effectiveness of trastuzumab in HER2 positive patients, some might suffer relapse with unknown reasons (3).

The current pathway-level understanding of cancer, though valuable, needs further investigation. Several computational methods have been developed to infer pathway activities for individual cancer sample (4, 5). For example, PARADIGM (4) adapted known pathways as Markov networks and inferred pathway activities from the observed multi-omics profiles with the EM algorithm. However, such methods usually relied on prior knowledge about signalling pathways. This may reduce the chance of obtaining novel biological insights, as our current knowledge is incomplete and biased towards well-studied pathways and genes (6). Furthermore, a recent study (7) has shown that databases of biological pathways are different from each other. Using different databases yielded disparate results in pathway analysis.

Therefore, computational researchers have proposed de novo methods (8–10) to identify latent patient pathways (factors) with multi-omics profiles. To ensure interpretability and computational feasibility, these methods assumed linear relationships among different types of omics data. Thus, it may not be sufficient to capture complicated biological relationships. On the other hand, some de novo methods were directly adopted from the machine learning community (11, 12), which may miss some biological features of cellular signalling pathways. For example, mutual exclusivity is a well-known phenomenon of somatic alterations in tumours, which is difficult to be captured by any machine learning model assuming additive effects. In fact, most methods (13–15) modelled mutual exclusivity with statistical tests.

In this work, we presented OR-gate Network (ORN), a de novo approach to infer the latent pathway activities in cancer. Fig. 1 shows how biological pathways (Fig. 1A) can be represented by an ORN (Fig. 1B). Although the order of signalling was ignored, ORN closely follows the causal mechanism from mutations to expressions. Each layer in ORN is connected by logical OR gates such that the output of a node is true if any input to the gate is true. This important property of the ORN agrees with the mutual exclusivity patterns observed in cancer genomics. On the other hand, when modelling the mechanism from latent pathway activities to transcriptomics, a pathway perturbed in a set of patients will cause differential expression of a set of genes. In this way, ORN generates the bicluster patterns a plethora of clustering algorithms aimed to recover in RNA expression data (16).

**Figure 1.**
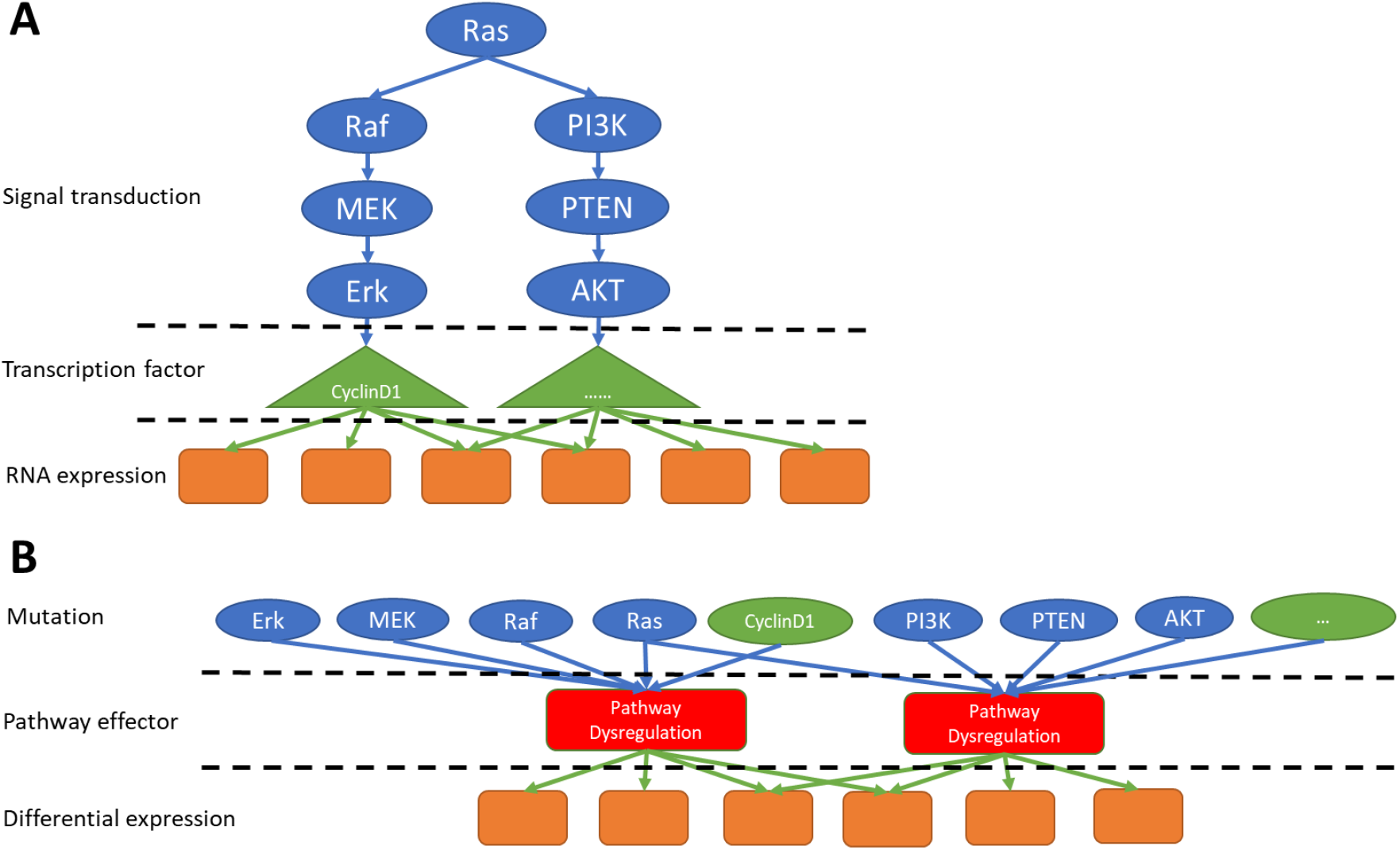
Figure A is the biologically plausible representation of signaling pathways. Given the difficulty in estimating the order of signal transduction, we replaced the realistic representation by introducing an additional variable indicating the status of pathway dysregulation, resulting in figure B. Genes in the genomic level are connected to pathway status with logical OR relationship. Pathway status is then connected to transcriptomics with the same logical OR relationships.

Since the generative process of ORN is consistent with both mutual exclusivity patterns in cancer genomics and bicluster patterns in cancer transcriptomics, we hypothesized that ORN can recover: (1) relations between pathways and downstream genes they regulate; (2) relations between SGA and the pathways they perturbed; (3) which pathway is dysregulated in which set of patients. To our knowledge, ORN is the first integrative model capable of inferring patient-specific pathway activities while soliciting corresponding SGAs and its regulon.

## MATERIAL AND METHODS

### Data preprocessing

Before applying ORN to real-word datasets, we need to transform genomic profiles and transcriptomic profiles into binary data. On the genomic side, we used two types of data: (1) non-silent gene-level simple somatic mutations (SNV) dataset; (2) gene-level copy number variation (CNV). An element in the CNV matrix was set to 1 if its original value was 2/-2. Any element with an absolute value less than 2 was set to 0. We then combined CNV and SNV data into a binary event matrix, that is, if the alteration of gene i was observed in either SNV or CNV in sample j, then the ijth element in the binary event matrix was set to 1.

For the transcriptomic profiles, we first remove genes with median expression counts lower than 10 across all samples. Then the Z scores provided by the CBioPortal platform were binarized. An element in the Z score matrix was set to 1 if its absolute value was greater than 1.96, otherwise 0.

To further filter the genes on the genomics side, we applied Multitask Lasso implemented in Scikit-learn (17). The genomic profiles were used as independent variables and the status of differential expression as targets. Somatic mutations with nonzero coefficients were retained as the input for our proposed algorithm.

### Model formulation

Before describing the full model, we need to first illustrate the key operation unit, leaky OR-gate. When there are several possible causes *X*_1_,*X*_2_, …, *X_n_* of an effect variable Y, the probability of Y should be:

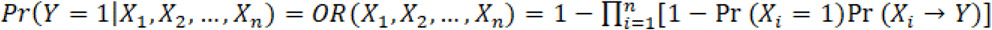

This is different from the definition in (18) with the introduction of (*X_i_* → *Y*). This term represents the probability that X causes Y. It allows us to infer the layered causal structure of ORN.

However, there are situations where **X** cannot fully capture the cause of Y. For example, pathway perturbation in cancer cells may be caused by methylation changes, not just somatic mutations. To deal with unmodeled causes, we introduced the leaky parameter, which is similar with the intercept in linear regression:

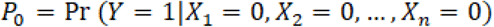

Hence the leaky OR gate function becomes:

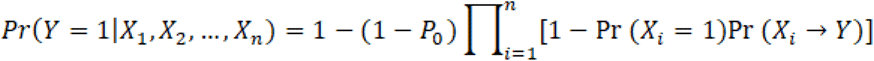

During implementation, *P*_0_ can be modelled by adding a row into the binary event matrix with all ones. For convenience, we will denote the vectorized OR gate function as *OR*(***X**, δ*) in all the following content, where X represents the vector [Pr (*X_i_* = 1)], and *δ* represents the vector [Pr (*X_i_* → *Y*)]. With the leaky OR-gate operation, we connected SGA to latent pathways and then to DEGs, as illustrated in Fig. 1B. After vectorization, the procedure of ORN is illustrated in Fig. 2. Let us denote S as the number of samples, M the number of genes in genomics, P the number of pathways, G the number of genes in transcriptomics. Let Mut be the S×M binary event matrix of somatic mutation, Path be the S×P pathway activity matrix, and Expr the S×G RNA expression matrix. Let U denote the M×P causal relationship matrix between SGA and pathways. Let Z denote the P×G causal relationship matrix between pathways and DEG. For the sth sample, the activity of the pth pathway is:

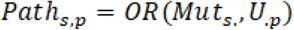

**Figure 2.**
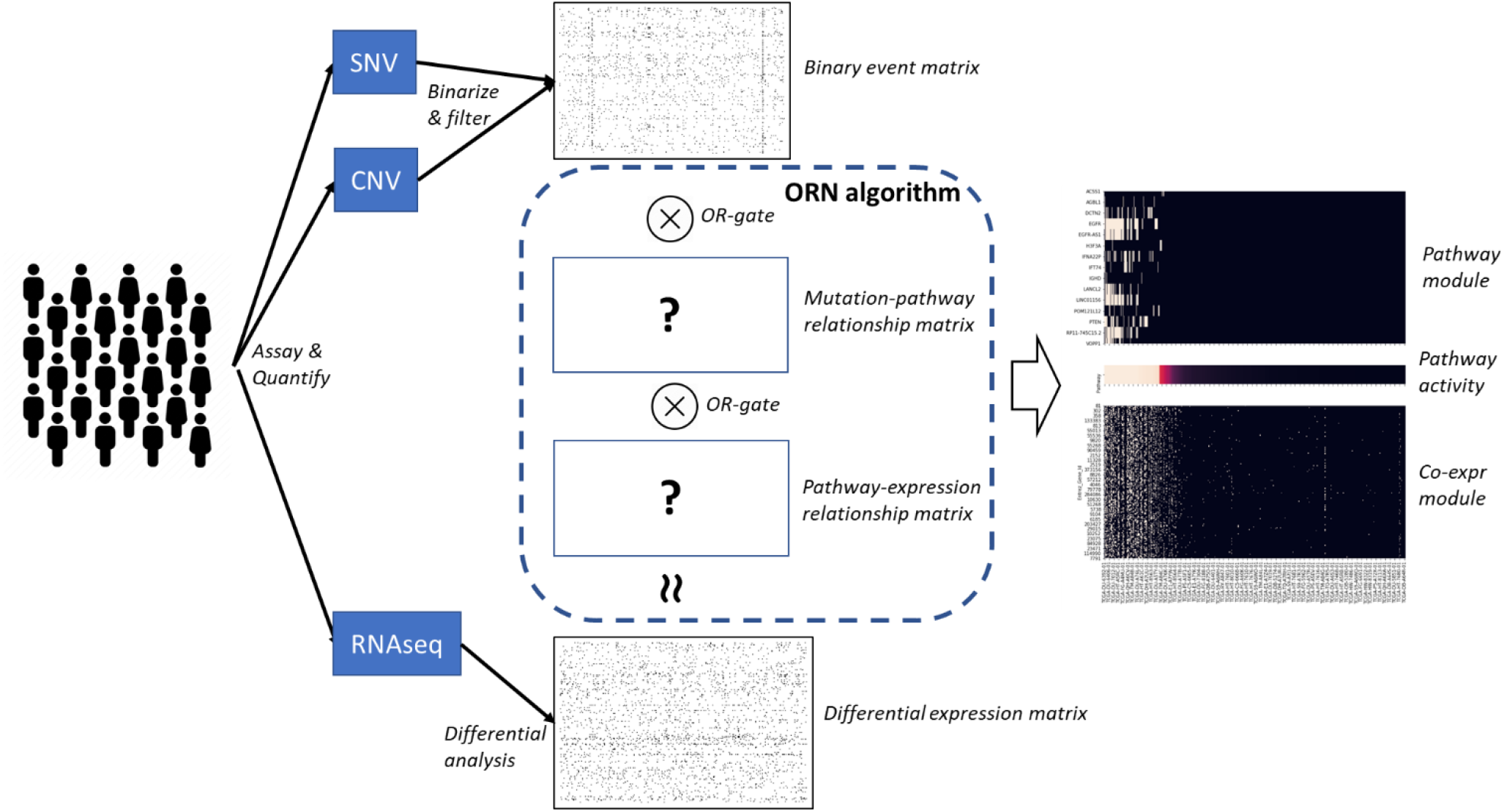
The source omics data for ORN consists of quantified matrices of single nucleotide variation (SNV), copy number variation (CNV), and gene expression (RNAseq). SNV and CNV were combined, binarized, and filtered on the genes’ side to produce a binary event matrix. As for RNAseq, we calculate robust Z score for each gene in each sample. We assumed Logical OR relations when binary events led to pathway dysregulation and, in turn, led to differential expression. ORN algorithm aimed to infer: (1) the relationship between somatic mutations and signaling pathways; (2) the relationship between signaling pathways and differential expressions. With the output from ORN, we can recover the latent pathways that were perturbed by somatic mutations and caused differential expression.

And the status of the *g*th DEG in the *s*th sample is:

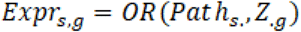

In this formulation, patient-specific pathway activities, *Path*, were represented as the logical units within the hidden layer.

The objective function to optimize for ORN is the overall log likelihood of the observed Expr given estimated Expr:

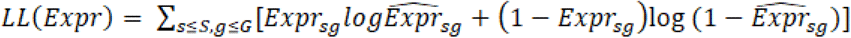

where 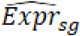 is the probability of differential expression estimated by the model, in other words, the last layer of ORN. Since *Path* and DEG can be computed by U and Z, they are the only parameters we need to estimate. Similar latent variable models, such as LDA (19), are usually computational expensive. However, we found that the layered structure of ORN is similar with the neural network architect. Thus, the powerful backward propagation approach, which is essential to all deep learning models, can be used to estimate ORN parameters. First, we need to reparameterize U and Z to *μ* and *ζ* such that the parameters are not bounded within [0,1]:

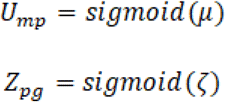

Then we updated *μ* and *ζ* during gradient descent. The gradient of *ζ* with respect to *LL(Expr)* is:

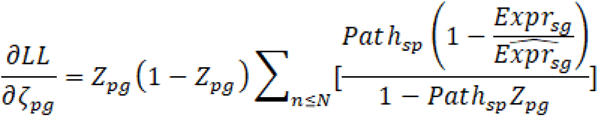

Using the chain rule, we can derive the gradient of *μ* with respect to *LL(Expr)*:

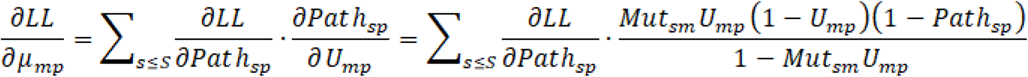

where the computation of *∂LL/∂Path_sp_* is symmetric to *∂LL/∂Path_sp_*.

To control the sparsity of parameters, we assumed U and Z are samples from the Beta distribution. Thus, the gradients above need to be modified. For example, the partial derivative of *ζ* should be modified as:

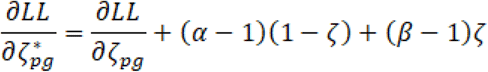

where *α* and *β* are the hyperparameters for Beta distribution. For all the experiments in this study, we set *α* = *α* = 0.95.

### Simulation and evaluation

Synthetic data was generated by following the noisy OR gate process. First, somatic mutations and the two relationship matrices were generated with Bernoulli distribution. Then *Path* and *DEG* were generated by performing noisy OR-gate computation with *P*_0_ = 0. To simulate the mutual exclusivity patterns observed in real data, we performed post pruning. That is, when several mutations belonging to the same pathway took place in the same sample, all but one of them were removed.

The artificial neural network (NN) was used as a baseline to evaluate ORN’s efficacy of inferring pathway activities. To ensure the neural network model is comparable with ORN, it had one hidden layer, and the activation function was sigmoid. In this way, the values of hidden neurons were also within [0,1]. We ran NN and ORN on 20 synthetic datasets and computed their reconstruction error and the average cosine similarity of pathways. Reconstruction error was computed as:

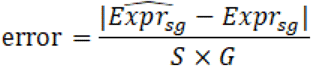

We further proposed Jaccard score, a more stringent criterion, to evaluate how ORN’s performance changes in various settings. To compute Jaccard score, we first need to compute Jaccard similarity for U and Z respectively:

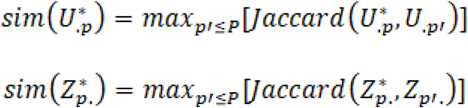

where U* and Z* are the true relationship matrix in synthetic data. Jaccard(A, B) is computed as:

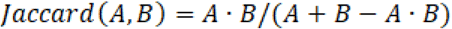

Then the Jaccard score for one dataset is:

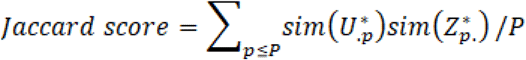

When applied to real world datasets, we extracted pathway activities together with its upstream SGAs and downstream regulon from the two relationship matrices. For example, if *U_mp_* < 0.5, then we concluded that mutation of gene m could disrupt pathway p. If *Z_pg_* > 0.1, then we concluded that the disruption of pathway p could cause differential expression of gene g. In this way, the set of genes affecting the same pathway were grouped into an upstream module, while the set of genes regulated by the same pathway were grouped into a downstream module. Note that in real data analysis, the cutoff for elements in Z was the top 5% value among all genes in the pathway p.

Since there is no ground truth for real data analysis, we run Gene Ontology (GO) enrichment analysis on the downstream modules to characterize the functional impacts of the latent pathways and its SGA modules.

### Public Data

We performed ORN on the high-throughput data of 514 lower grade glioma patients. It was downloaded from The Cancer Genome Atlas (TCGA). After preprocessing, 511 samples remained with 15501 DEGs and 670 SGAs. The number of latent pathways was set to 10.

To evaluate how different cancers may have perturbed the same pathways, we also applied ORN to the METABRIC dataset (20) downloaded from CBioPortal. It contained 2173 samples of somatic variant profiles and 1904 samples of RNA-seq. After preprocessing, we retained a binary event matrix with 1092 genes and a binary DEG matrix with 24256 genes. Both matrices had 1866 samples. ORN was trained with 15 latent pathways.

## RESULTS

### ORN was effective in recovering OR-gate relationships

Synthetic data were generated according to the generative process described in Methods. The number of pathways was set to 5; The number of samples, SGAs, and DEGs were all set to 1000. This was referred to as the standard setting.

We proposed Jaccard score and reconstruction error to evaluate the performance. Jaccard score can measure the concordance between inferred relationship matrices and the ground truth. Details of calculation was described in Methods.

From the standard setting, each condition was changed separately to see how they affected the performance. As shown in Fig. 3, ORN has achieved almost perfect recovery (>99%) in the standard setting. However, ORN’s performance dropped over 20% when the number of samples dropped from 1000 to 300, or the number of mutations increased from 1000 to 3000 mutations. Note that when the number of DEGs were reduced to 500, the performance remained the same.

**Figure 3.**
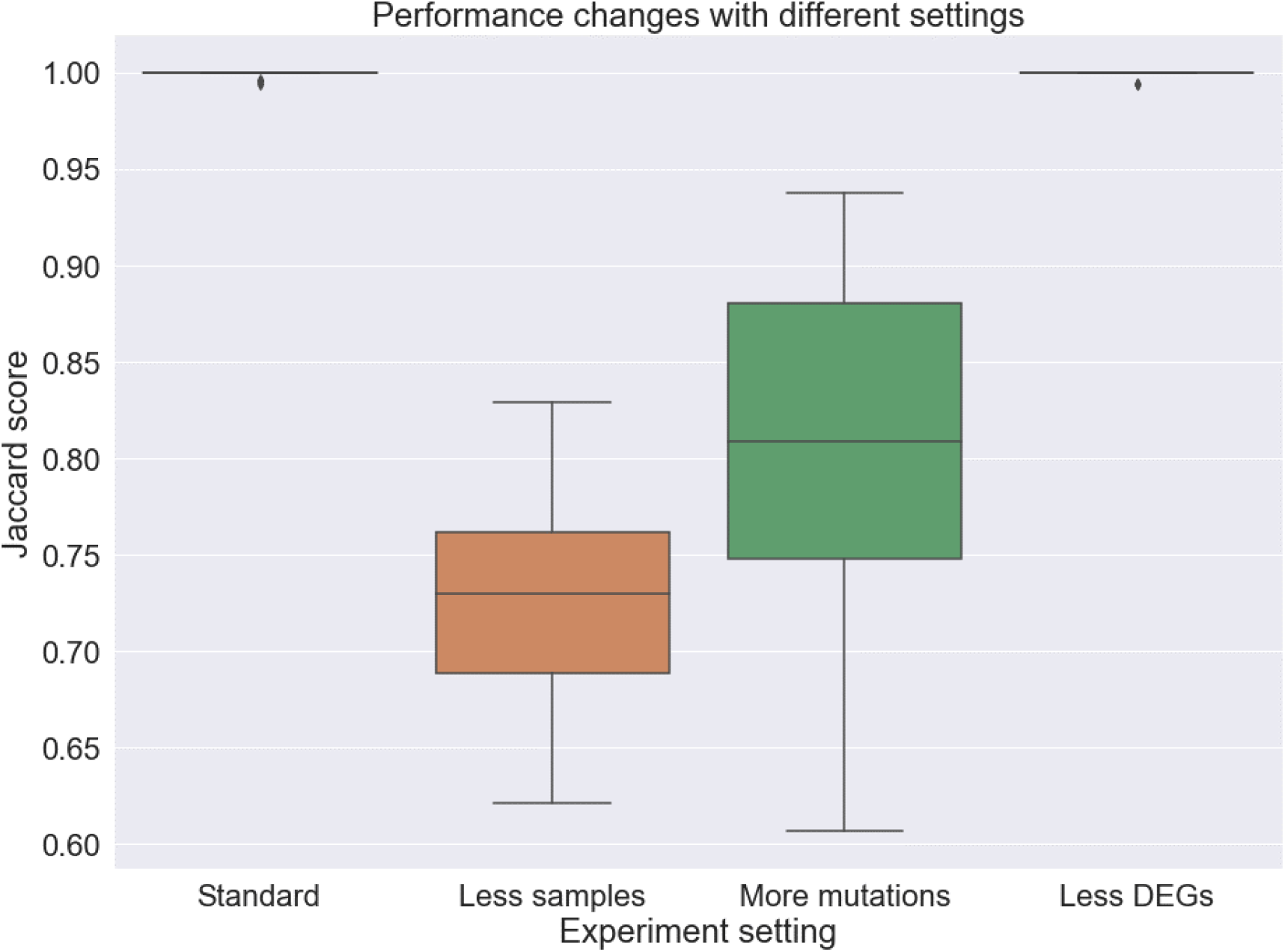
In the standard setting, ORN has recovered latent pathways with almost perfect accuracy. When the number of samples decreased to 300 or the number of mutations increase to 3000, the median Jaccard score has decreased to 73% and 81% respectively. Reducing the number of DEGs did not affect the performance of ORN.

### ORN provided more insights than neural network

We cannot identify similar algorithms that only used high-throughput data to infer pathway activities. However, we found that artificial neural networks (NN) with sigmoid activation function can also produce binary values in the hidden layer. In addition, both ORN and NN used backward propagation to optimize parameters. Thus, we designed a neural network architecture similar with ORN and used it as a baseline.

In the synthetic experiment, although NN converged to comparable reconstruction error as ORN, its accuracy was in pathway recovery was only around 50%. This showed that NN is less capable of capturing all the signals in the data.

Similar results were observed when we applied NN to the glioma dataset (described in the next section). The relationship matrix estimated with NN is much more redundant than ORN. GO enrichment analysis of the downstream modules (see Supp. Table 1) also showed that NN could only capture less than 5 major aspects with 10 hidden neurons, while ORN can cover different biological aspects of glioma with each latent pathway. As shown in Fig. 5, the relationship matrix learned by NN contains much redundancy, while each pathway in ORN regulated different sets of genes with few overlaps. This indicated that the biological mechanism from somatic mutations to transcriptomic profiles could be more accurately characterized by the OR-gate logic imposed by ORN rather than conventional non-linear relationships.

**Figure 4.**
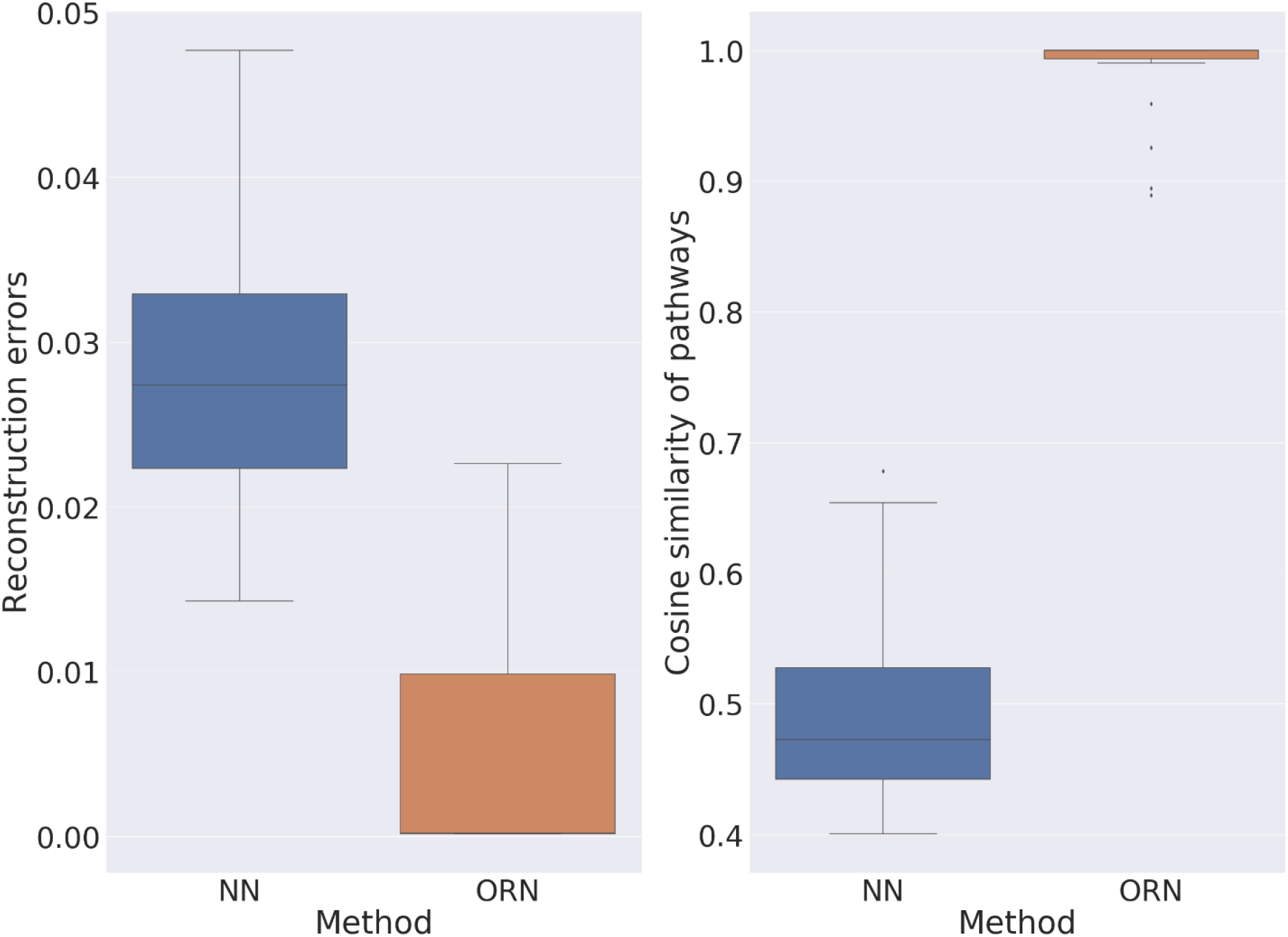
Boxplot on the left showed the distribution of prediction error of NN and ORN across 20 synthetic experiments. Boxplot on the right showed the distribution of cosine similarity between the inferred pathways and ground truth.

**Figure 5.**
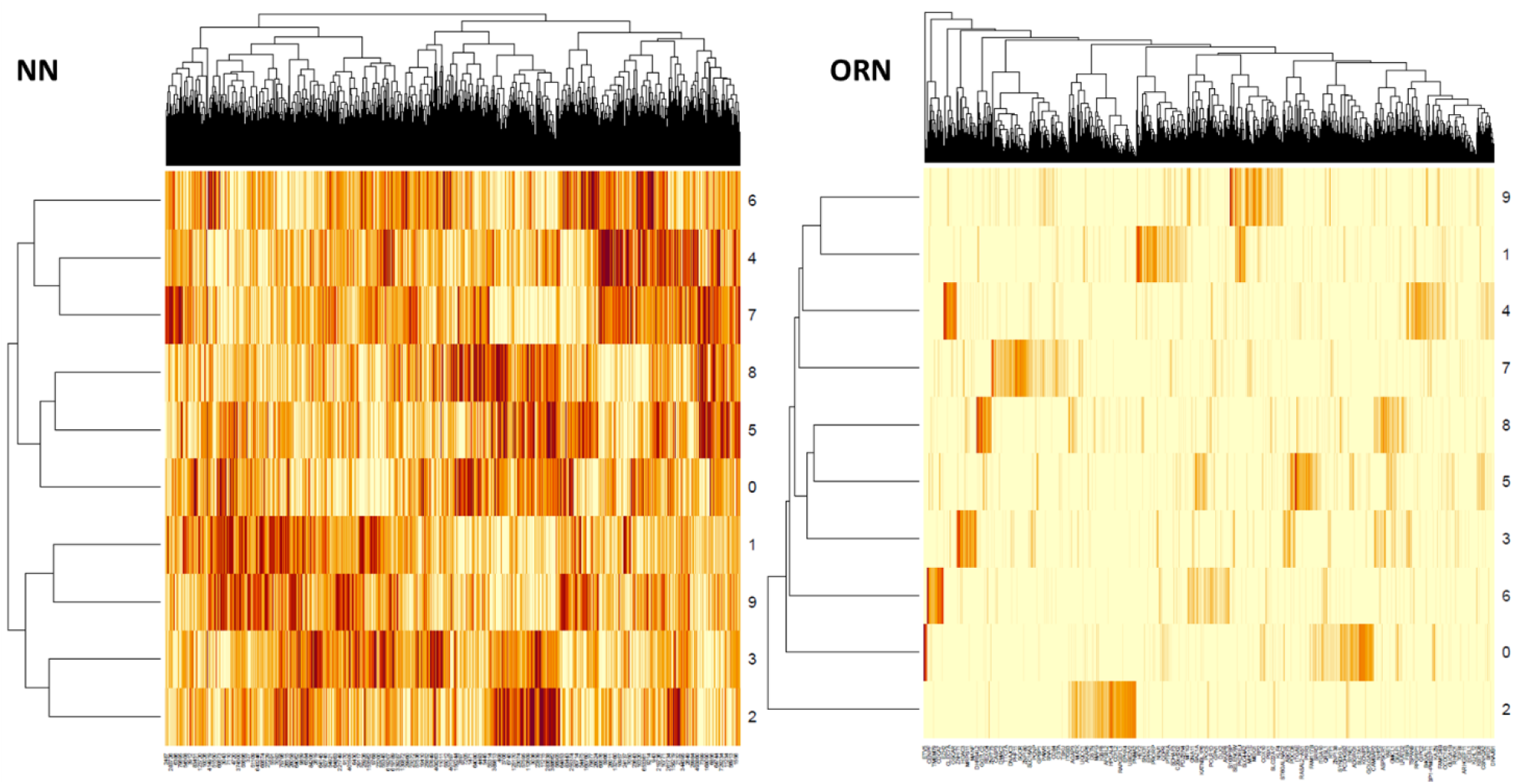
Heatmap representation of the relationship matrix between pathways and differential expression after row normalization. Relationship matrix generated by a neural network (NN) contains many redundant signals, while ORN automatically pushes for sparsity. Each latent pathway in ORN has uniquely caused a subset of genes to express differentially.

### ORN detected pathways closely related to patient survival

After applying ORN to the lower grade glioma dataset. Pathway activities showed that pathway 6 and pathway 7 had significant impacts on patient survival (Fig 6). We performed Gene Ontology (GO) enrichment analysis on the top DEGs in these pathways (see supplement table 2). The downstream module of pathway 6 is mostly related to DNA processing activities. Cancer samples with this pathway dysregulated probably have compromised genomic instability (21), leading to worse survival. Its upstream module includes CDK13 (22), H3F3A (23), IDH1 (24), PTEN (25), SNRPE (26) that are closely related to DNA repair or DNA replication.

**Figure 6.**
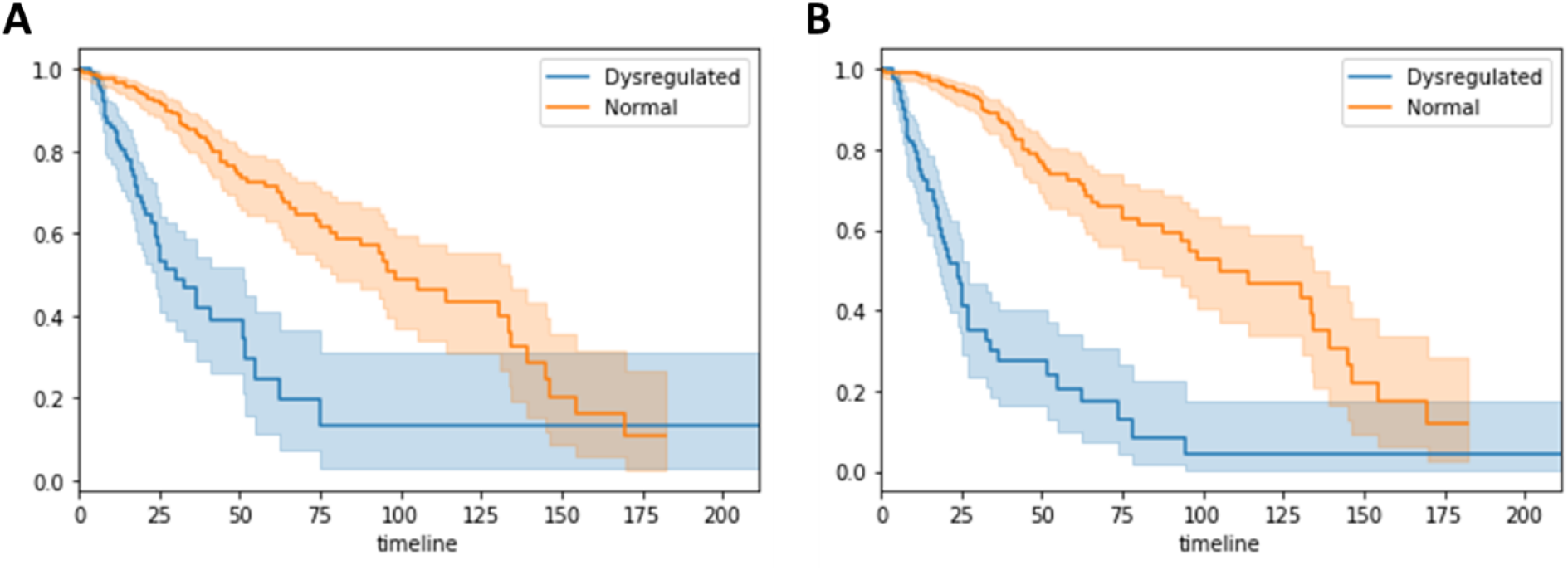
LGG patients with pathway 6 (A) and pathway 7 (B) dysregulated have worse overall survival. X-axis is in the unit of month; Y-axis represents the proportion of each subgroup. 102 patients’ pathway 6 were dysregulated, 100 patients’ pathway 7 were dysregulated. Both groups have 62 samples in common.

As for pathway 7, we found that PTEN, H3F3A, and POM121L12 were shared by the upstream modules in both pathways. However, the top 300 DEGs caused by the two pathways have no genes in common. GO enrichment analysis showed that downstream modules are related to neutrophil activities, Ras signal transduction, and viral genome replication. We conjectured that cancer samples with pathway 7 dysregulated exhibited viral infection and its immune response. Since virus infection can drive glioma formation (27), This subgroup of patients may be more likely to progress to malignancy and worse survival.

### ORN characterized common mechanisms in cancer

As for the results in the METABRIC dataset, upstream modules of six latent pathways were almost identical and hence merged into one union (pathway 0, 1, 2, 7, 9, 14 in supplement table 3). This module contained well-known oncogenes such as TP53, PTEN, PIK3CA, and MAP3K1. The corresponding pathway was dysregulated across all samples (probability above 0.5). This pathway likely represented the common cause of breast cancer. GO enrichment analysis (see supplement table 3) showed that its corresponding downstream module mostly involved in mitosis such as G1/S transition of mitotic cell cycle phase transition (GO:0044772). Pathway 5 also exhibited similar downstream effects but had different SGAs. Most notably, MIR604 was in the upstream modules of pathways. Studies have shown that polymorphism of MIR604 was related to the development of hepatocellular carcinoma (28) and the metastasis of colorectal cancer (29). MIR604 was differentially expressed in breast cancer (30). Still, to our knowledge, the impact of MIR604 mutation in breast cancer has not been investigated.

In the glioma dataset, we found pathway 4 to be closely related to immune response. Downstream modules of pathway 4 were enriched for various immune response, including cytokine-mediated signalling and toll signalling. This subgroup of patients may not be responsive to immune therapy. Within this pathway was the mutation of interferon alpha 21 (IFNA21), which played an important role in inflammatory response and toll signalling. IFNs were also identified as major factors of patient response to various cancer therapies (31). Moreover, we found that PTK6 and SRMS within the same upstream module. The product of two genes work closely together as intracellular kinases (32) and promotes invasive prostate cancer (33). However, they are rarely studied in the context of glioma and immune response.

Like glioma, one particular pathway in breast cancer captured abnormal immune response in a subgroup of cancer samples. The downstream module in pathway 3 is related to immune response, including T cell activation (GO:0042110), regulation of immune response (GO:00507006), inflammatory response (GO:0006954). The upstream module included CDC20, COLEC12, MED8, MPL, SOX5, and OTUD1. CDC20 was known to be related to T cell activation. COLEC12’s protein product is associated with innate immunity (34). SOX5 was shown to be related to B cell proliferation (35). Another interesting gene was MED8. Studies showed that MED8 was important to regulate resistance against bacteria in plants (36). Meanwhile, MED8 was implicated in renal cell carcinoma. However, it is rarely investigated in the case of breast cancer and innate immunity. As for OTUD1, a recent study (37) has shown that its induction by RNA virus may inhibit innate immune response.

### ORN detected pathway dysregulation specific to lower grade glioma

Although not related to patient survival, other pathways in glioma samples also captured different aspects of molecular characteristics. For example, pathway 0 is closely related to the biosynthesis of cholesterol, steroid, and alcohol, while cholesterol metabolism has recently been studied as a potential therapeutic target (38). In addition, downstream modules of pathway 0 contained differentially expressed genes enriched for central nervous system development. In the corresponding upstream module, we identified SZT2 (39) and TIAM1 (40) to be closely related to nervous system development. Other mutations, such as CPAMD8 and RUBP1, exhibited mutual exclusivity and similar expression patterns. Yet these two genes have not been studied in terms of central nervous system development.

As for the breast cancer samples, the upstream module of pathway 8 contained several well-known driver mutations, including KRAS, APC, and ARID1A. As shown in Fig 7, most genes in this upstream module exhibited mutual exclusivity, while these mutations caused differential expression of a similar set of genes. The downstream module was enriched for telomere and t-circle formation, which were well-known factors for cancer initiation and tumour survival (41). In the upstream module, the relation between telomere and APC (42), KRAS (43), ARID1A (44), PRKG1 (45) were reported. Although mutually exclusive to the four genes above, we have not found research linking BAP1, MIR604, and MICAL3 to telomere activities.

**Figure 7.**
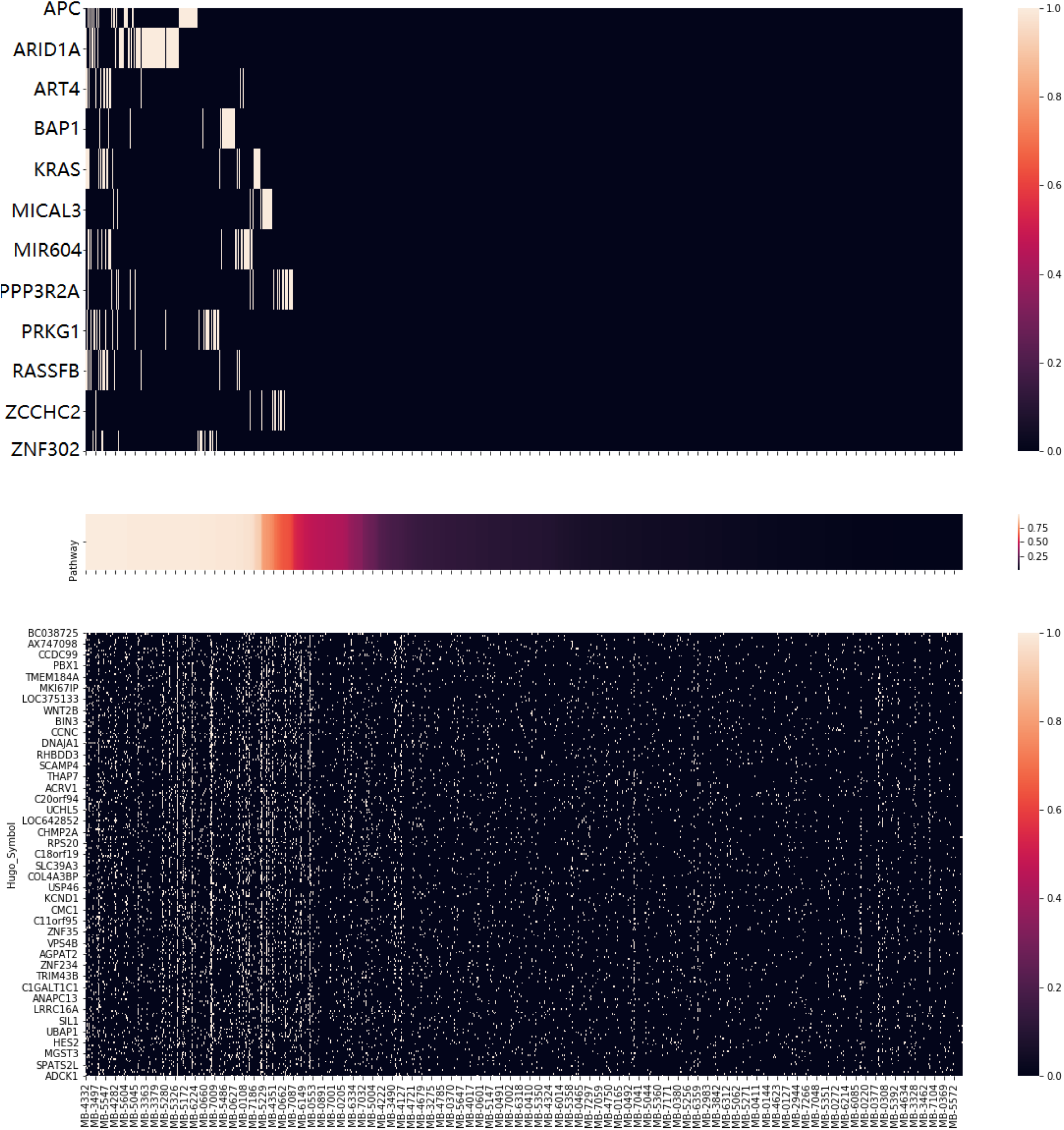
Illustration of pathway 8 in breast cancer samples. Cancer samples sorted by pathway activities (middle figure) was the X-axis shared by the three subplots. The upper figure showed the mutation event of the upstream module (cutoff 0.5), while the bottom figure showed the heatmap of differential expression of the downstream module (roughly top 1% related). The top figure showed patterns of mutual exclusivity, while the bottom showed a strong correlation between pathway activities and differential expression.

## DISCUSSION

In this study, we proposed ORN, the first de novo method to infer latent pathway activities from genomic profiles and transcriptomic profiles of cancer patients. Compared with the traditional neural network method, ORN provided much more insights into how the somatic mutations function together. It also implied that the OR-gate logic assumed in ORN is suitable to the mechanism from mutations to differential expressions.

In the real data analysis with lower grade glioma and breast cancer, ORN has recovered major mechanisms consistent with current knowledge, such as abnormal DNA repair ability and immune response. Glioma patients with these dysregulated pathways had lower survival rates. ORN further revealed mechanisms specific to cancer types, such as the steroid metabolism and nervous system development in glioma. We identified several somatic mutations that might be related to certain malfunctions in cancer cells, worthy of further biological investigation. However, ORN requires in-depth analysis to obtain useful insights. In the future, we will try to develop statistical tests to automatically return meaningful genes in both upstream and downstream modules.

For the METABRIC dataset, none of the latent pathways generated by ORN is significantly related to patient survival. We conjectured that the reason was two-fold: (1) Somatic mutations may not be the only source of variance of RNA expression. Sharma, et al showed that copy number alterations, epigenetic changes, transcription factors, and microRNAs collectively explain, on average, only 31– 38% and 18–26% expression variation (46); (2) compared with glioma samples, cellular constitution in breast cancer samples was probably more diverse. To handle the first issue, we need to include more data sources in a principled way. In the future, epigenetic profiles may be included to inform the coregulation of RNA expression. To deal with the second issue, future research needs to incorporate reliable complete deconvolution algorithms in the data preprocessing step.

For some genes in the upstream module, we failed to find evidence supporting their associations to the downstream module. Although some of them were likely to provide novel molecular insights, many were false positive. Upon closer investigation, we believed there are two major sources of false positives: (1) passenger mutations that exclusively occur in highly mutated samples. For example, ACSS1 was in most upstream modules of glioma because it only occurred in highly mutated samples, which had most pathways dysregulated. (2) passenger mutations within the same copy number variation event as driver mutations. When a set of genes mutated in almost the same set of patients, it is likely that only one of them contributed to the pathway dysregulation.

During analysis, we also found that different upstream modules may share a subset of somatic mutations. For example, pathway 6 and pathway 7 in glioma have PTEN in common. Thus, it is possible that hierarchical structures of SGA functions can be inferred from the overlapping upstream modules. In the future, we may provide more convenient visualization utilities to analyse the hierarchies among upstream modules.

We proposed ORN to infer latent pathway activities from high-throughput profiles of cancer samples. Application of ORN in lower grade glioma and breast cancer detected latent pathways closely related to patient survival. ORN also connected somatic mutations to key mechanisms of cancer, such as DNA repair and innate immune response. Although some mutations’ function (e.g., MIR604) was not supported by literature, they were mutually exclusive to well-known driver mutations and caused differential expression in a similar subset of genes. We encouraged biological researchers to use ORN to infer personalized pathway activities and generate novel hypotheses for targeted therapy.

## DATA AVAILABILITY

Processed data from METABRIC can be downloaded at: https://www.cbioportal.org/study/summary?id=brcametabric

Processed data of TCGA-LLG can be downloaded at: https://xenabrowser.net/datapages/?cohort=TCGA%20Lower%20Grade%20Glioma%20(LGG)

Python implementation of ORN is available at: https://github.com/LifanLiang/ORN

