## Supplemental Figures for "ORN: Extracting Latent Pathway Activities in Cancer with OR-gate Network"

June 5, 2020

#### 1 TCGA lower grade glioma (LLG) pathway visualization

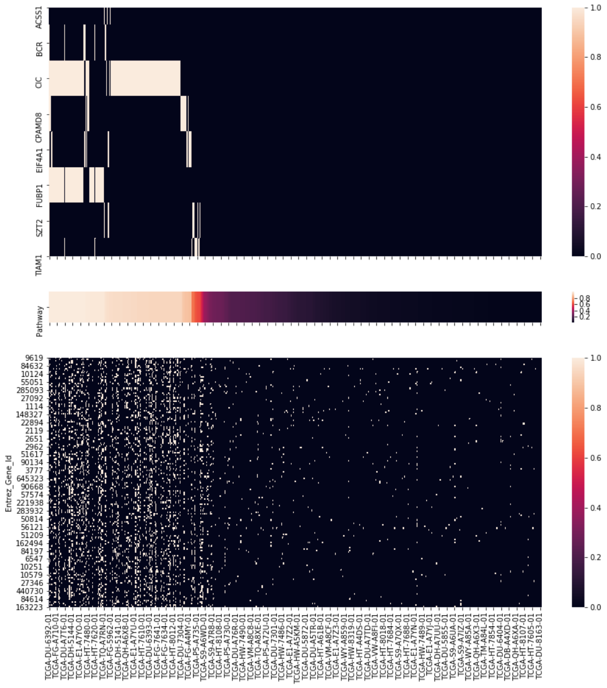


Figure 1: Visualization of Pathway 0 in lower grade glioma

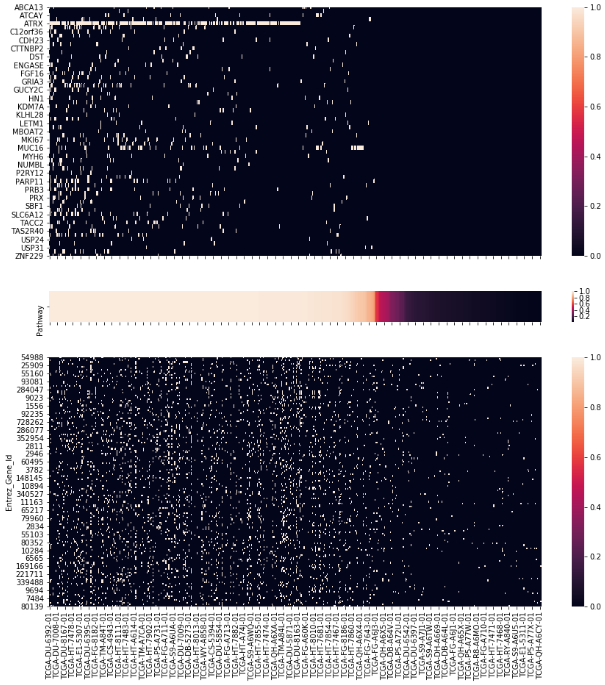


Figure 2: Visualization of Pathway 1 in lower grade glioma

### 2 METABRIC pathway visualization

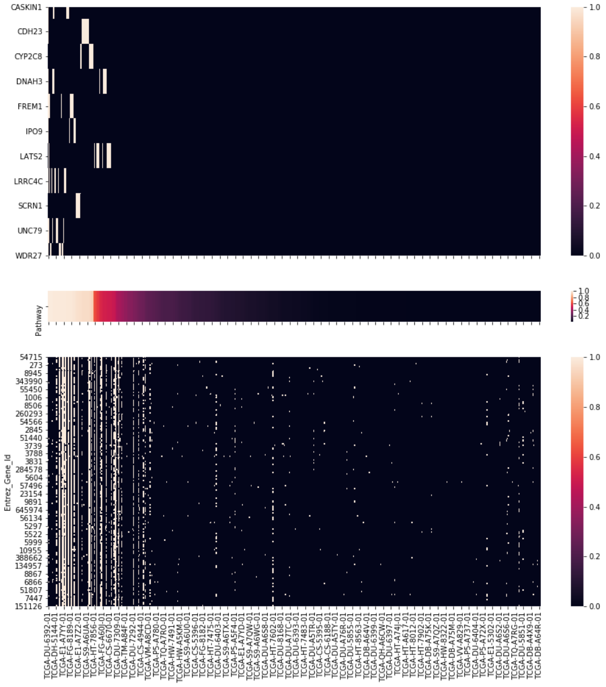


Figure 3: Visualization of Pathway 2 in lower grade glioma

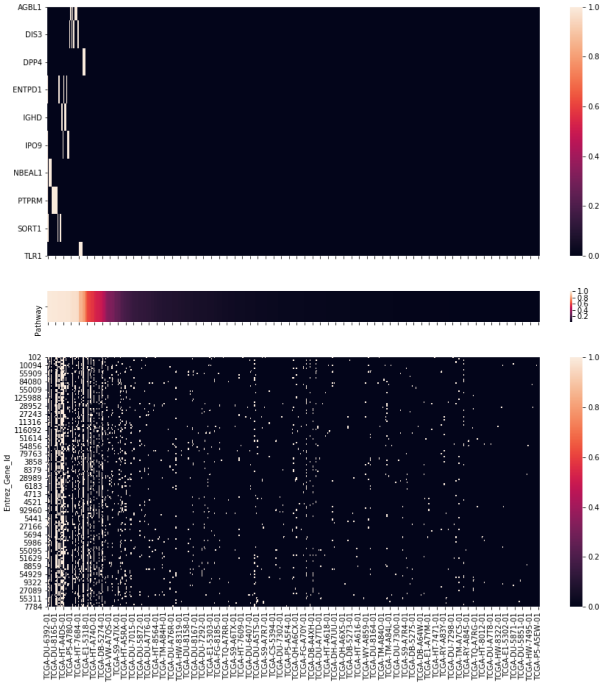


Figure 4: Visualization of Pathway 3 in lower grade glioma

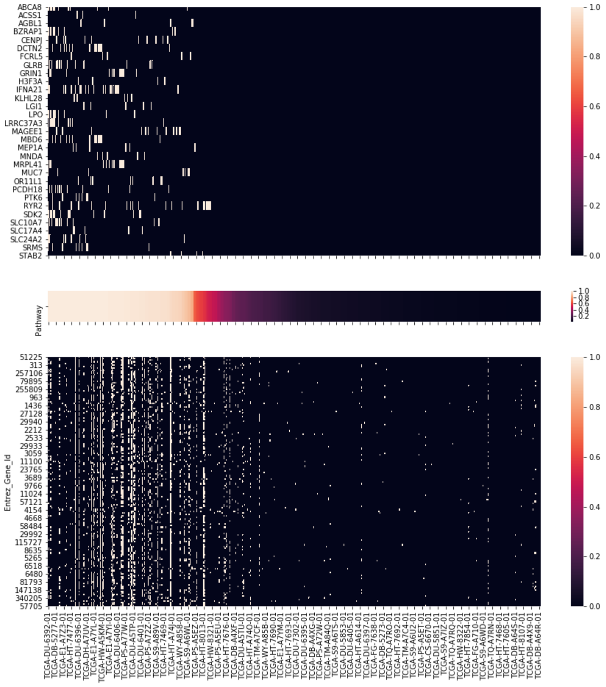

Figure 5: Visualization of Pathway 4 in lower grade glioma

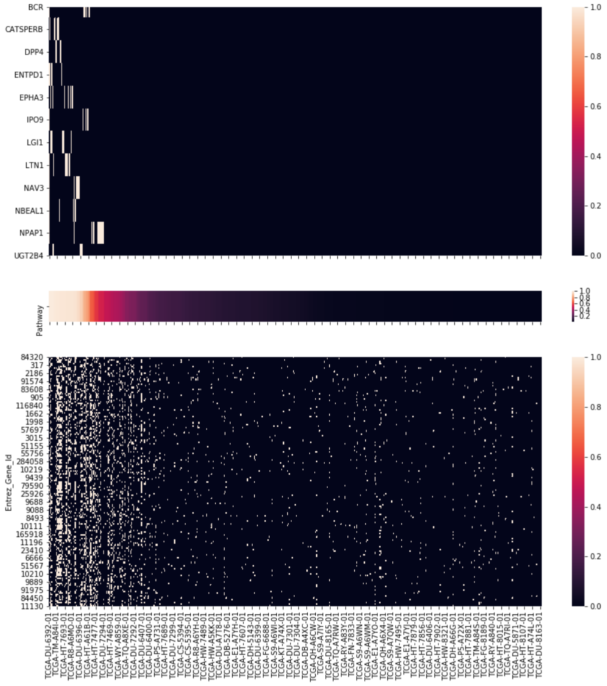


Figure 6: Visualization of Pathway 5 in lower grade glioma

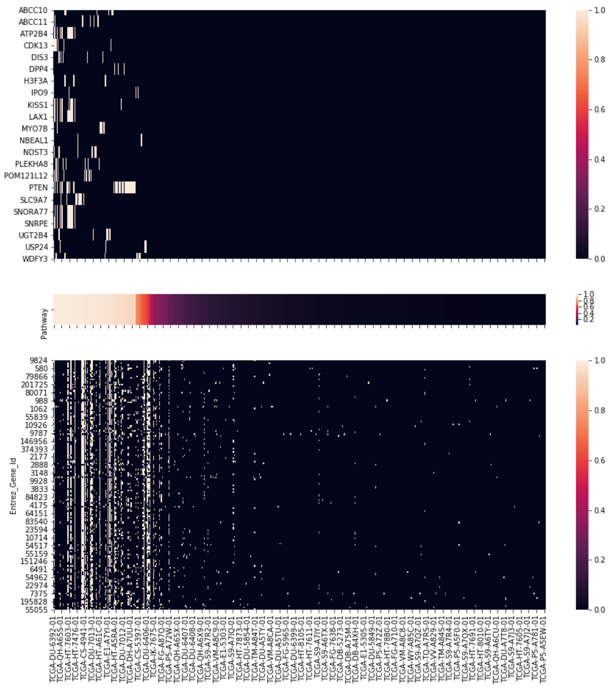


Figure 7: Visualization of Pathway 6 in lower grade glioma

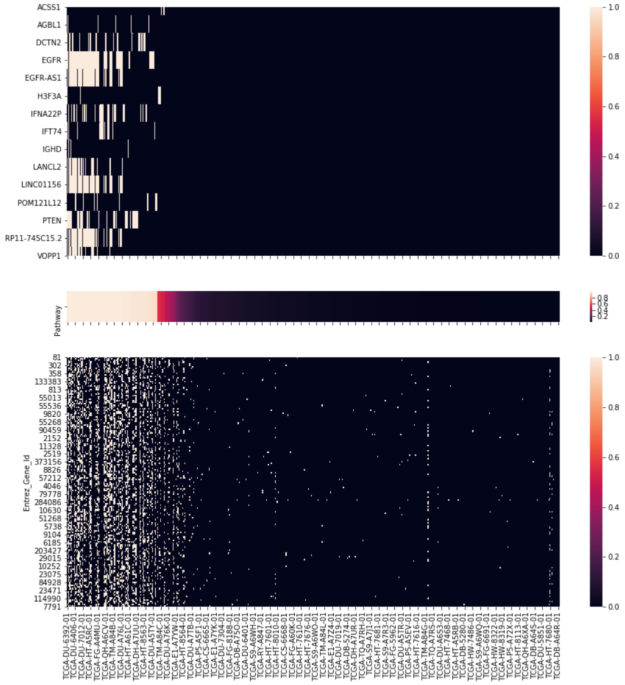

Figure 8: Visualization of Pathway 7 in lower grade glioma

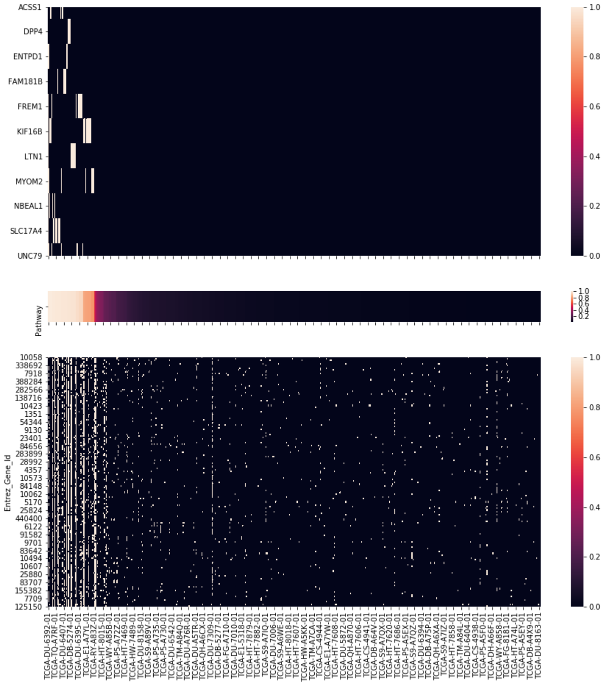

Figure 9: Visualization of Pathway 8 in lower grade glioma

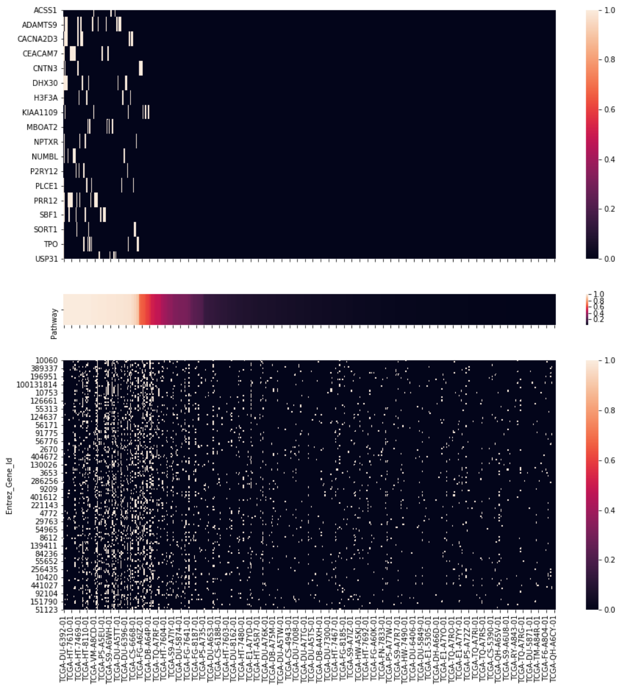

Figure 10: Visualization of Pathway 9 in lower grade glioma

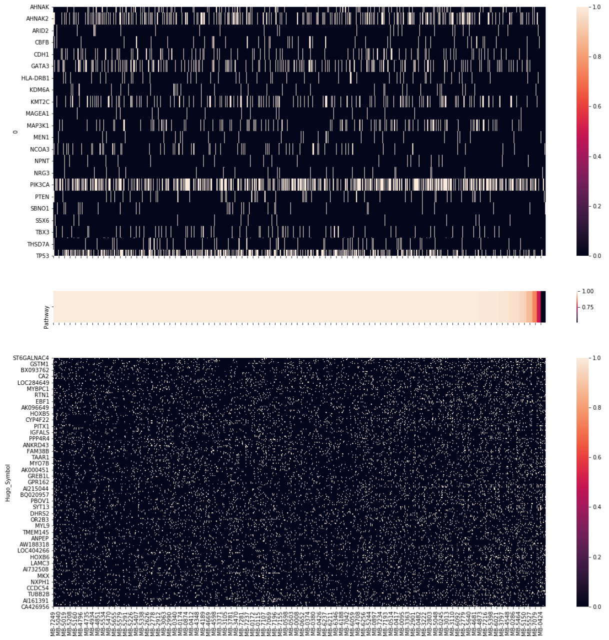


Figure 11: Visualization of Pathway 0 in Metabric

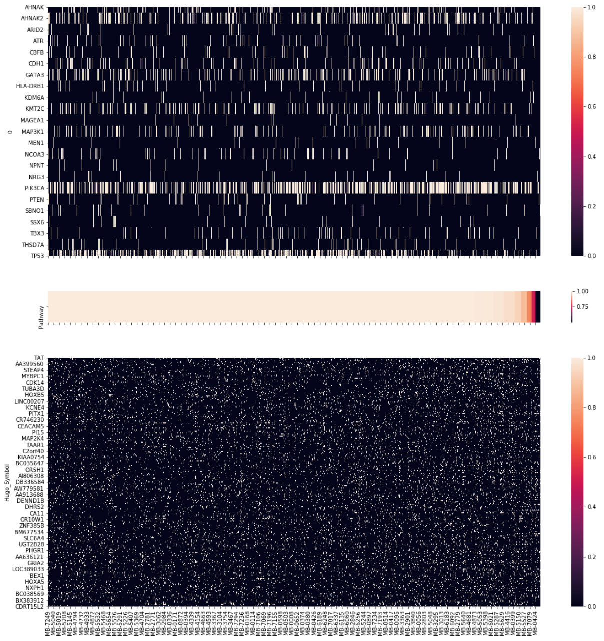


Figure 12: Visualization of Pathway 1 in Metabric

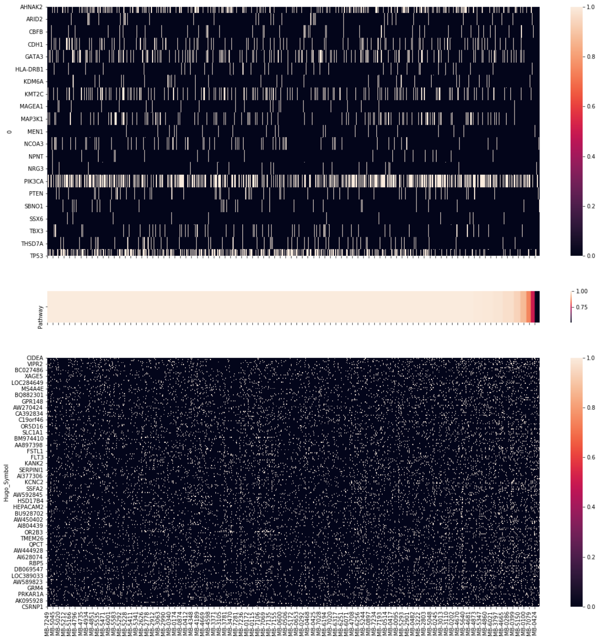


Figure 13: Visualization of Pathway 2 in Metabric

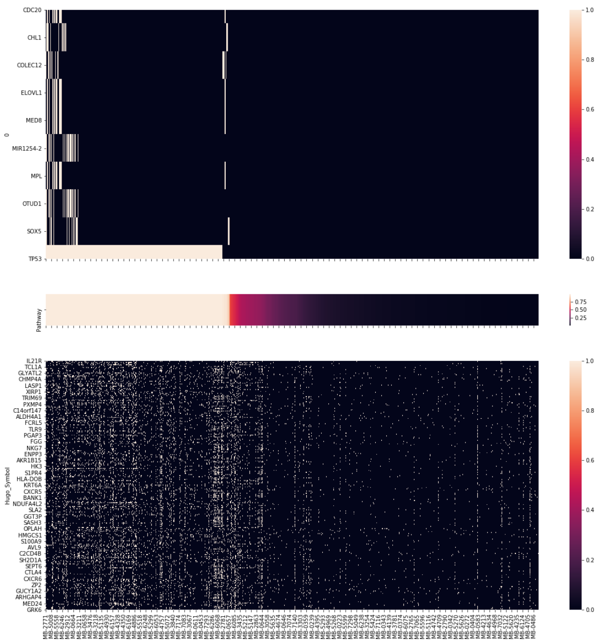


Figure 14: Visualization of Pathway 3 in Metabric

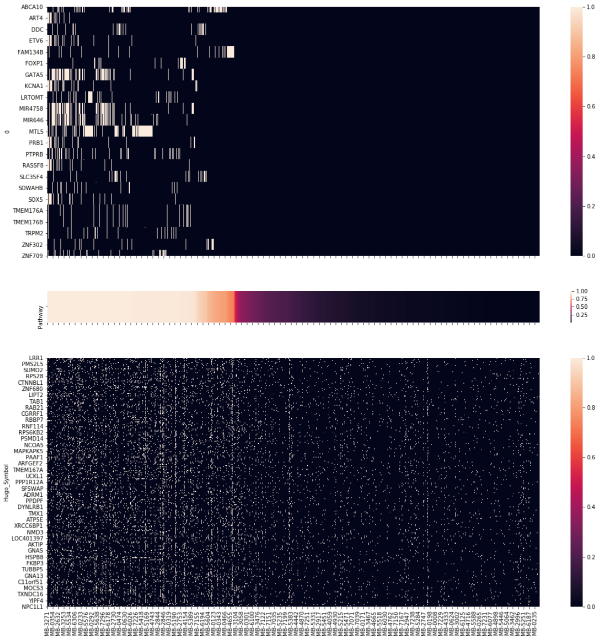


Figure 15: Visualization of Pathway 4 in Metabric

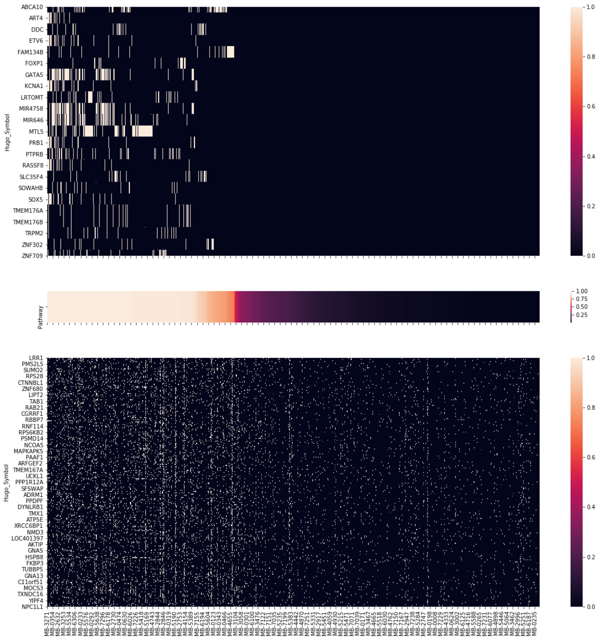


Figure 16: Visualization of Pathway 6 in Metabric

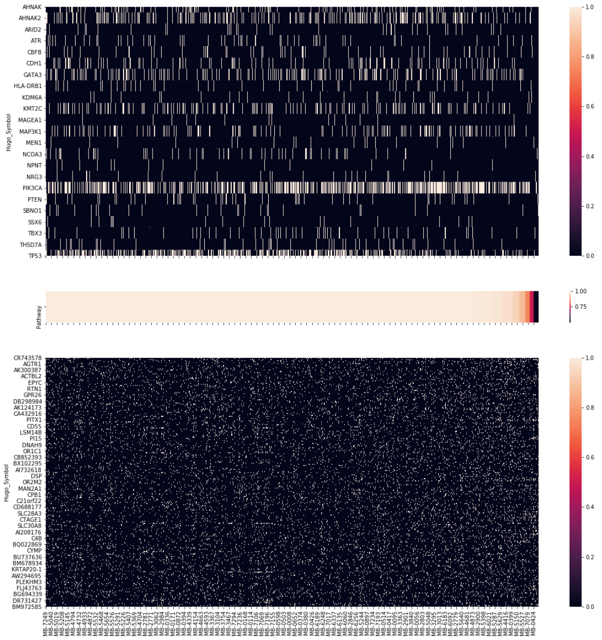


Figure 17: Visualization of Pathway 7 in Metabric

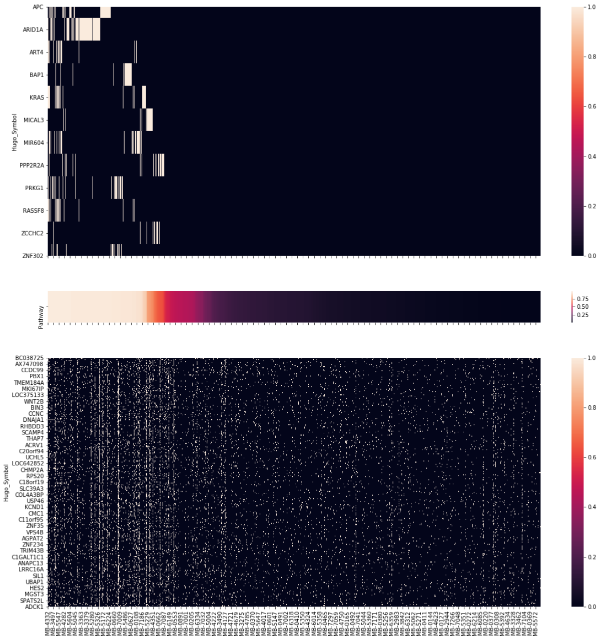


Figure 18: Visualization of Pathway 8 in Metabric

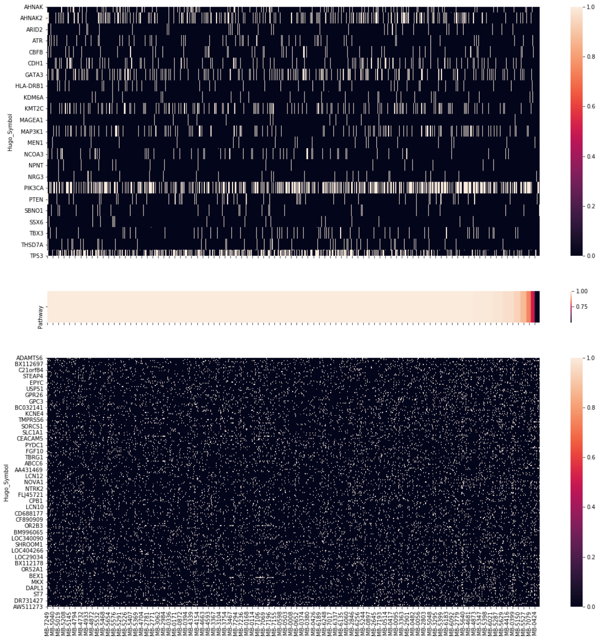


Figure 19: Visualization of Pathway 9 in Metabric

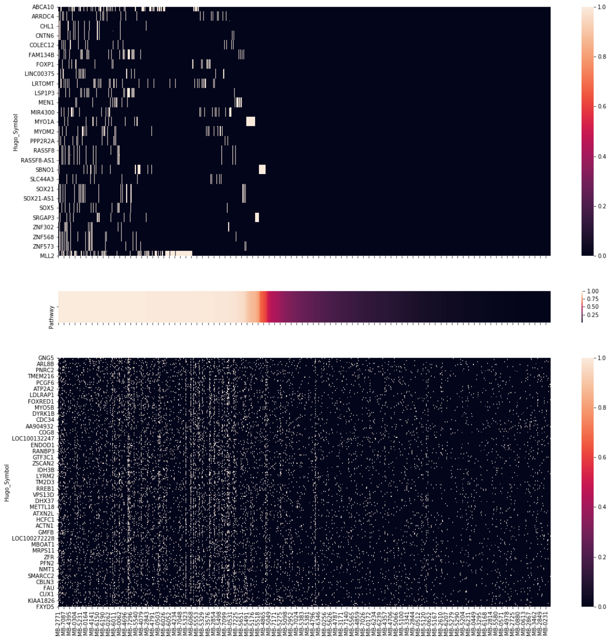


Figure 20: Visualization of Pathway 10 in Metabric

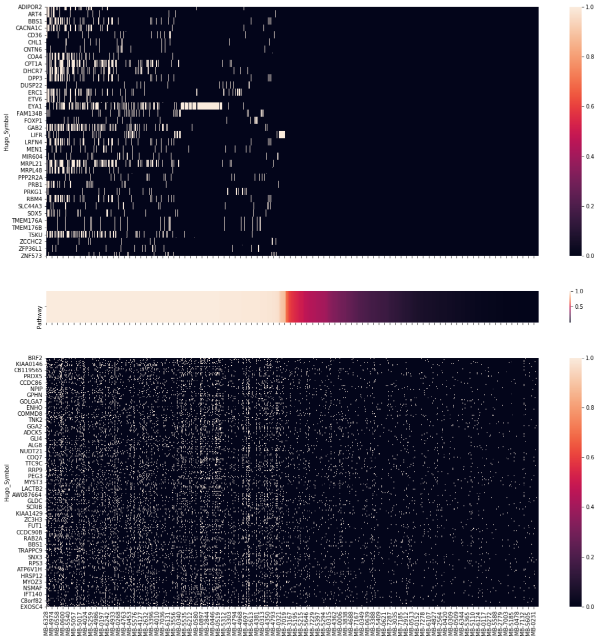

Figure 21: Visualization of Pathway 11 in Metabric

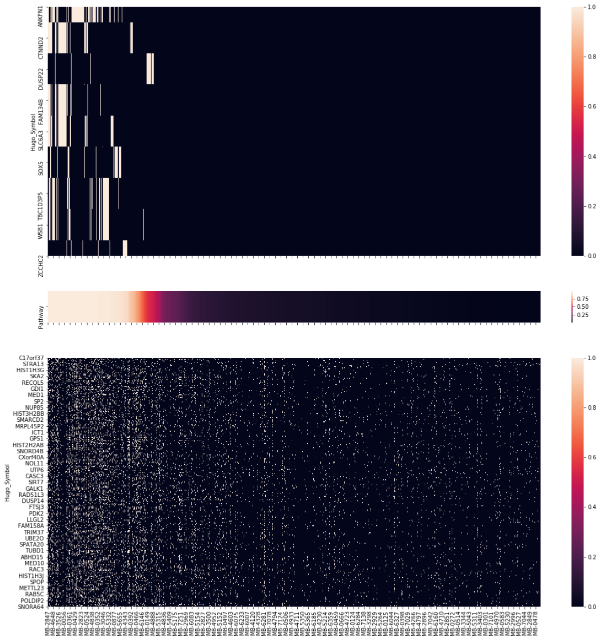

Figure 22: Visualization of Pathway 12 in Metabric

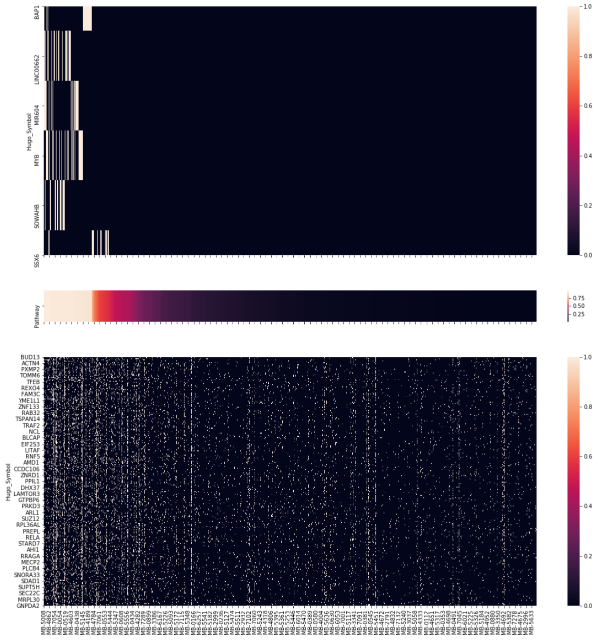


Figure 23: Visualization of Pathway 13 in Metabric

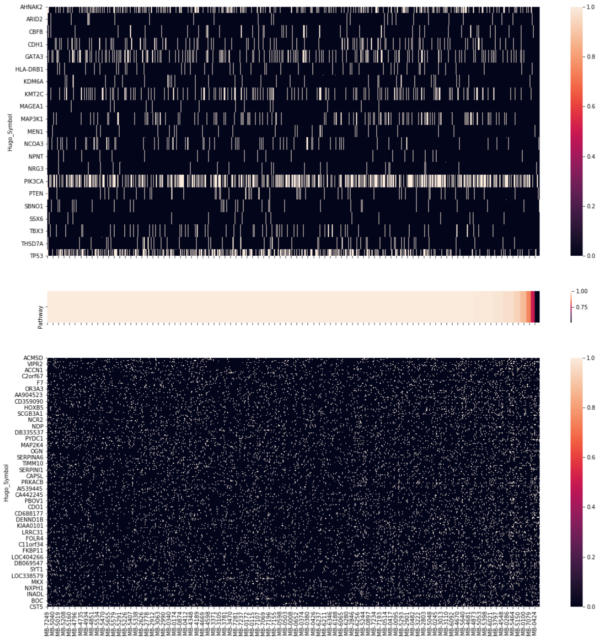


Figure 24: Visualization of Pathway 14 in Metabric
